# Characterization of *Staphylococcus lugdunensis* biofilm reveals key differences according to clonal lineage and iron availability

**DOI:** 10.1101/2025.07.25.666850

**Authors:** Laurie Destruel, Sandrine Dahyot, Laurent Coquet, Magalie Barreau, Stéphanie Legris, Marie Leoz, Maxime Grand, Xavier Argemi, Gilles Prevost, Nicolas Nalpas, Emmanuelle Dé, Sylvie Chevalier, Martine Pestel-Caron

## Abstract

To understand the mechanisms involved in the evolutionary success of *Staphyloccocus lugdunensis* clones, we compared the biofilm-forming ability of representative strains of the seven clonal complexes (CCs) in rich and iron-restricted conditions, and characterized the extracellular matrix (ECM) of two highly biofilm-forming strains under each condition.

Over 90% of the 49 *S. lugdunensis* strains produced biofilm in both conditions, with a level of production depending on the iron availability and clonal lineage. Two behaviors were observed: a significantly higher production in rich medium than in iron-restricted medium for CC1, CC2, and some CC3 strains, and the opposite phenomenon for CC6 ones. Analysis of the ECM of two representative strains using confocal microscopy showed that biofilm of the CC3 strain in rich medium contained similar amounts of proteins, eDNA and polysaccharides while that of CC6 strain was predominantly proteinaceous. Under iron-restricted conditions, biofilm structure and composition of both strains completely differed from those obtained in rich conditions. The proteomic analysis of their biofilm ECM by liquid chromatography coupled to tandem mass spectrometry identified 321 proteins common to both strains, mainly intracellular and in particular ribosomal. Of note, 202 proteins differed between the strains in terms of abundance, with a higher proportion of membrane proteins in the CC3 strain.

This study performed on a large cohort of strains shows that *S. lugdunensis* biofilm-forming capacity is strongly associated with CC and iron availability. This analysis of biofilm-associated proteins in *S. lugdunensis* opens the way to propose new molecular targets for anti-biofilm strategies.

**IMPORTANCE:** The ability of *S. lugdunensis* to produce biofilm is considered as a critical virulence factor. As biofilm is strongly associated with persistence and difficult-to-treat infections, characterizing biofilm production and composition, particularly in iron-deficient environments encountered during infection, can provide a better understanding of therapeutic failures. Our work is the first to be carried out on such a large collection of *S. lugdunensis* clinical strains. It shows that this species is a strong biofilm producer, even in an iron-deficient environment, and that the composition of its matrix varies according to both genetic background of the strain and environmental conditions. Moreover, investigating the biofilms protein matrix of two *S. lugdunensis* strains provides insights into identification of potential targets for biofilm eradication.

## INTRODUCTION

*Staphylococcus lugdunensis* is a particularly virulent species among coagulase-negative *Staphylococci* (CoNS), with a clinical pathogenic potential quite similar to that of *Staphylococcus aureus* (1). *S. lugdunensis* can cause a wide range of infections, including skin and soft tissue infections, bone and joint infections, catheter-related infections, and particularly aggressive and destructive infective endocarditis (2). To monitor the epidemiology of these infections, typing methods have been developed, such as the multilocus sequence typing (MLST), which defines sequence types (STs) grouped into clonal complexes (CCs) (3, 4). Worldwide, seven CCs have been described, and interestingly, CC1 and CC3 were mainly identified in hospital settings, both in Asia and in France (4–7).

To date, *S. lugdunensis* pathogenesis remains unclear, with few well-characterized virulence factors (1, 2, 8). Biofilm-forming ability appears as a characteristic of the species, and seems to be of particular importance in bone and joint infections (notably prosthetic joint infections) (9, 10). Whole genome sequencing analyses have identified the presence of the *ica* locus in all the 21 *S. lugdunensis* genomes belonging to various CCs (11). The *icaADBC* locus encodes enzymes involved into biosynthesis of the poly-N-acetylglucosamine (PNAG) polysaccharide, which is commonly constitutive of the biofilm matrix of other CoNS and *S. aureus* strains (12). However, through biofilm dispersion assays, confocal laser scanning microscopy (CLSM) or immuno dot blot analyses, *in vitro* studies initially failed to detect the presence of PNAG in *S. lugdunensis* strains and revealed that biofilm matrix was mainly composed of proteins (13, 14). Nevertheless, recent studies using the same methodological approaches have shown that polysaccharides along with proteins were the major components of *S. lugdunensis* mature 24h-biofilm (15, 16). However, these studies were conducted on small collections of strains (11 to 38) not phylogenetically characterized, and biofilm analyses were carried out under laboratory optimal growth conditions in rich media (9, 10, 13, 16–19). Only two studies have investigated biofilm formation under *in vivo*-like conditions, by analyzing biofilm-forming capacity of up to nine *S. lugdunensis* strains (clinical strains of unknown CC) in iron-deficient conditions (20, 21). They revealed that the biofilm matrix comprised proteins and not PNAG under these low-iron growth conditions, and demonstrated the particular role of the iron acquisition system (Isd) proteins in promoting biofilm in this restricted environment. Recently, Cho *et al.* have characterized for the first time the extracellular matrix (ECM) proteome of a *S. lugdunensis* strain (NCCP 15630, Korea, CC unknown) in rich medium in comparison with planktonic condition, by liquid chromatography coupled to tandem mass spectrometry (LC-MS/MS) (22). Interestingly, among proteins identified only in biofilm were proteins involved in iron transport, like ferrous iron transport B and the probable heme-iron transport system permease protein IsdF, as well as a DUF5084 domain-containing protein (22).

In this context, we first sought to further explore the role of iron-restricted conditions in biofilm production in a large collection of *S. lugdunensis* phylogenetically characterized strains. Second, we investigated the biofilm composition by CLSM as well as the ECM proteome by LC-MS/MS of two strong biofilm-producing strains belonging to different CCs.

## MATERIALS AND METHODS

### Bacterial strains

Forty-nine *S. lugdunensis* strains were included in this study: 12 carriage strains and 37 responsible for human infections ***(Supplementary Table 1)*** (3, 4, 23). Forty-eight strains originated from diverse French University Hospitals (Bordeaux, Lyon, Montpellier, Nancy, Nantes, Rouen, Strasbourg, Tours) and one from Sweden (Kronoberg County). Strains were previously characterized by MLST (24) and/or *fbl*-typing (4). Forty-five strains belonged to 7 CCs (11 CC1, 5 CC2, 11 CC3, 4 CC4, 5 CC5, 5 CC6, and 4 CC7) and 4 were singleton STs (1 ST13 and 2 ST28) ***(Supplementary Table 1)***. *S. epidermidis* ATCC 35984 (RP62A) strain was used as a biofilm producing control strain (25).

### Growth curve assays

Growth kinetics of the 49 *S. lugdunensis* strains were performed in rich medium (trypticase soy broth, TSB, Bio-Rad, Marnes-la-Coquette, France) and in iron-depleted medium (RPMI 1640, Sigma-Aldrich, Saint-Louis, Missouri, USA) with 1% of casamino acids (Sigma-Aldrich) (RPMI). Overnight cultures were incubated in TSB or RPMI under shaking (150 rpm) at 37°C and then diluted to OD_600_ of 0.01. Aliquots (200 µL per well) of standardized samples were inoculated in triplicate in a 96-well polystyrene microtiter plate. OD_600_ measurement was performed every 15 min with the *Spark®* multimode microplate reader (Tecan, Männedorf, Switzerland) over 24h at 37°C under continuous double-orbital agitation (108 rpm). Data were obtained from biological triplicates. Generation time and maximal growth rate were determined with the R^©^ software (v.3.6.0). Statistical analyses were performed by Kruskall-Wallis and post hoc Dunn’s test on GraphPad Prism^©^ software.

### Biofilm formation

Biofilm formation was assessed by a crystal violet staining assay (adapted from Stepanovic *et al.* 2007) (25) in TSB supplemented with 1% glucose (TSBG) and in RPMI for all the 49 *S. lugdunensis* strains and the control producing strain *S. epidermidis* ATCC 35984 (25). Overnight cultures were diluted to obtain an OD_600_ of 0.01. Microtiter plates (96-wells) were filled in triplicate with 200 µL of bacterial suspension and incubated for 24h at 37°C under static conditions. Supernatants were removed and wells were washed once with sterile water. Remaining adherent cells were stained for 20 min with 200 µL of 0.5% crystal violet (Sigma-Aldrich) at room temperature, then washed twice with sterile water, and crystal violet was solubilized by ethanol 96% for 20 min. OD_590_ was measured with the *Spark®* multimode microplate reader (Tecan). For each plate, three wells contained medium alone (blank). Four biological replicates were performed. Statistical analyses were performed by Wilcoxon Mann Whitney test on R^©^ software.

For each strain, the average OD_590_ values for all strains were compared to the cut-off value (ODc). ODc was calculated using the following formula: ODc = average OD ^blank^ + 3 x standard deviation^blank^. Strains were classified as non-producer (OD_590_ ≤ ODc), weak biofilm producer (ODc < OD_590_ ≤ 2 x ODc), moderate producer (2 x ODc < OD_590_ ≤ 4 x ODc) and strong producer (4 x ODc < OD_590_) as previously described by Stepanovic *et al.* (25).

### Biofilm inhibition assays

Assays were performed for a CC3 (03MC) and a CC6 (SL-55) strain using a method adapted from Panda and Singh study (26). The 24h-biofilm was grown in TSBG in 24-wells microplates. Solutions composed of proteinase K (100 µg/mL, Sigma-Aldrich), sodium metaperiodate (100 µg/mL, Sigma-Aldrich) or DNAse I (100 µg/mL, Sigma-Aldrich) were added onset of the culture to determine the amounts of proteins, exopolysaccharides and eDNA, respectively. Wells without enzymatic or chemical treatment were used as controls. Wells were then treated and stained with 0.5% crystal violet as described above. Data were obtained from three biological replicates. Results were expressed as ratio between mean of treated wells and mean of control wells. Statistical analyses were performed by Wilcoxon Mann Whitney test on R^©^ software.

### Confocal Laser Scanning Microscopy (CSLM)

A 48h-biofilm of the CC3 (03MC) and CC6 (SL-55) strains were grown in TSBG and RPMI in 24-well glass bottomed microplates (Greiner bio-one, Les Ulis, France). Briefly, the initial OD_600_ was adjusted to 0.08. After 48h of incubation at 37°C, cells were washed with 300 µL of 0.9% NaCl to remove planktonic cells. The membrane integrity of the cells embedded in the biofilm was evaluated by LIVE/DEAD *Bac*Light Bacterial Viability Kit^®^ (Invitrogen, Waltham, Massachusetts, USA) according to the manufacturer’ instructions. Cells were stained by adding the green fluorescent probe Syto-9 (50 nM, Invitrogen). Biofilm matrix components were labeled using the red fluorescent SYPRO Ruby dye (Thermo Fisher Scientific, Waltham, Massachusetts, USA) for proteins, the red fluorescent 1,3-dichloro-7-hydroxy-9,9-dimethyl-2(9H)-acridinone (DDAO) (1 µM, Euromedex, Souffelweyersheim, France) for eDNA, and the blue fluorescent Calcofluor White (200 µg/mL, Sigma-Aldrich) for β1-3 and β1-4 exopolysaccharides. The CLSM observations of biofilms were performed using a Zeiss LSM710 confocal microscope (Carl Zeiss Microscopy, Oberkochen, Germany) using a x63 oil immersion objective. Images were taken every 0.5 µM. For visualization and processing of 3D images, the Zen 2.1 SP1 zen software (Carl Zeiss Microscopy) was used. Data were obtained from three biological replicates. Quantitative analysis of images was performed with COMSTAT software (https://www.comstat.dk) (27, 28).

### ECM proteins characterization

#### Matrix extraction

ECM proteins of the CC3 (03MC) and CC6 (SL-55) strains were extracted according to an adaptation of the protocol of Chiba and coll. (29). Overnight cultures were diluted to obtain OD_600_ of 0.01. Briefly, the biofilm was grown in 100 mL of TSBG for 24h in an Erlenmeyer flask containing 2g of glass wool under agitation (30 rpm) at 37°C. Glass wool was rinsed in 100 mL of sterile water. The ECM was extracted with 15 mL of 1.5 M NaCl. After gently shaking for 5 min on a roller shaker, bacterial cells were removed by centrifugation (5000 g for 15 min) and filtered through a 0.22 µM filter. Then, the ECM was precipitated overnight at 4°C in three volumes of 96% ethanol. Finally, the ECM was concentrated by centrifugation (9000 g for 30 min at 4°C), washed in 70% ethanol, air-dried and resuspended in water. The experiments were performed in biological quadruplicate for each strain.

#### Trypsin digestion and nano LC-MS/MS

ECM extracts (25 μg) were incubated overnight at 37°C with dispersin B (20 ng/µL) and loaded on SDS-PAGE gel (5% polyacrylamide) and migrated for 45 min at 20 mA. Proteins were stained with Coomassie blue. A single band containing all the extracted proteins was excised for endoprotease digestion carried out with an automated system (MultiPROBE II, PerkinElmer, Waltham, Massachusetts, USA). The protein bands were washed several times in water, dehydrated with acetonitrile, dried, incubated in a reductive solution (dithiothreitol 5 mM) and alkylated in iodoacetamide 25 mM. Then, the trypsin digestion was performed overnight with 1 µg of trypsin (Promega, Madison, Wisconsin, USA). The resulting peptides were recovered from the gel by incubation twice for 15 min in acetonitrile and once for 15 min in a 0.1% TFA (TriFluoroacetic Acid) solution. Peptides were dried, concentrated in 0.1% TFA solution and quantified by colorimetric peptide assay (Pierce Quantitative Peptide Assays, Thermo Fisher Scientific).

For mass spectrometry analysis, 0.2 µg of protein digests were injected in an LTQ Orbitrap Elite mass spectrometer (Thermo Fisher Scientific) equipped with a nano-ESI source coupled to a nanoliquid chromatography (Easy-nLC II, Thermo Fisher Scientific). Peptides were separated by using a reversed phase C18 column (NikkyoTechnos, Japan) with a linear gradient of 15% to 55% of B (mobile phase A: water/0.1% TFA and phase B: Acetonitrile/0.1% TFA) over 120 min. The mass spectrometer was operated in data dependent mode to automatically switch between Orbitrap-MS (from m/z 300 to 2000) and LTQ-MS/MS acquisition.

#### Protein identification and quantification

Mass spectrometry data (raw data files) were processed using Progenesis QI software (Waters, Nonlinear Dynamics). Briefly, after peptide map alignment and normalization, the analysis of variance (ANOVA) with statistic filters was performed to select peptides showing significant and reproducible difference expression levels. When the *p-*value was < 0.05, the associated MS/MS spectra were exported for peptide identification with Mascot (Matrix Science v2.6.0) against the NCBI database restricted to *S. lugdunensis* HKU09-01 (RefSeq assembly: GCF_000025085.1). The identification searches were performed with variable modifications for oxidation of methionines, carbamidomethylation of cysteines, pyro-glutamate (Q and E) and with a maximum of 2 missed cleavages. MS/MS spectra were searched with a mass tolerance of 5 ppm for precursor ions and 0.35 Da for MS/MS fragments. Only peptides exhibiting significant Mascot individual ion score were retained. The proteins abundance was calculated by summing the abundances of all associated identified peptides. We first selected proteins identified from at least 2 peptides, and second compared the abundance of proteins using an ANOVA with a *p-*value < 0.05 and a power > 0.8 between the two strains. In addition, only the proteins showing a 1.8-fold ratio between the two strains was considered as significantly changing in abundance.

Proteins’ cellular locations was predicted using the NCBI database and the PSORT website (http://www.psort.org/psortb). Participation in biological functional pathways was determined using the Kyoto Encyclopedia of Genes and Genomes (KEGG; http://www.genome.jp/kegg).

## RESULTS

### Iron restriction affects *S. lugdunensis* biofilm formation according to the phylogenetic lineage

Growth assays were performed on 49 *S. lugdunensis* strains in TSB and RPMI to evaluate the potential impact of phylogenetic lineage on growth in rich and iron-restricted conditions. Mean generation time and maximal growth rate in TSB were 37.3 min ± 4.1 and 1.1 ± 0.1, respectively, and 80.7 min ± 33.1 and 0.6 ± 0.2 in RPMI. The generation time was 2.2 times higher and the maximum growth rate was 1.9 times lower in iron-restricted medium compared to TSB. No growth difference was observed according to the strains or the CCs, whatever the medium. In TSB, the final stationary OD_600_ was 1.2 ± 0.4 while it was lower in RPMI (0.5 ± 0.2). Although strain-dependent, the maximum OD_600_ did not seem to depend on the CC of the strains.

We then searched for a correlation between clonal lineages and ability to form biofilm and investigated whether experimental condition of iron limitation was correlated to a greater ability to produce biofilm. In TSBG, almost all strains (47/49, 95.9%) produced biofilm, among which 40.8% (*n* = 20) were strong-, 32.7% (*n* = 16) moderate- and 22.5% (*n* = 11) weak-biofilm producers (***Figure 1A***). The two strains that did not produce biofilm belonged to CC5. CC2 strains produced significantly (*p*<0.05) more biofilm than strains of all other CC (***Figure 2***).

**Figure 1.**
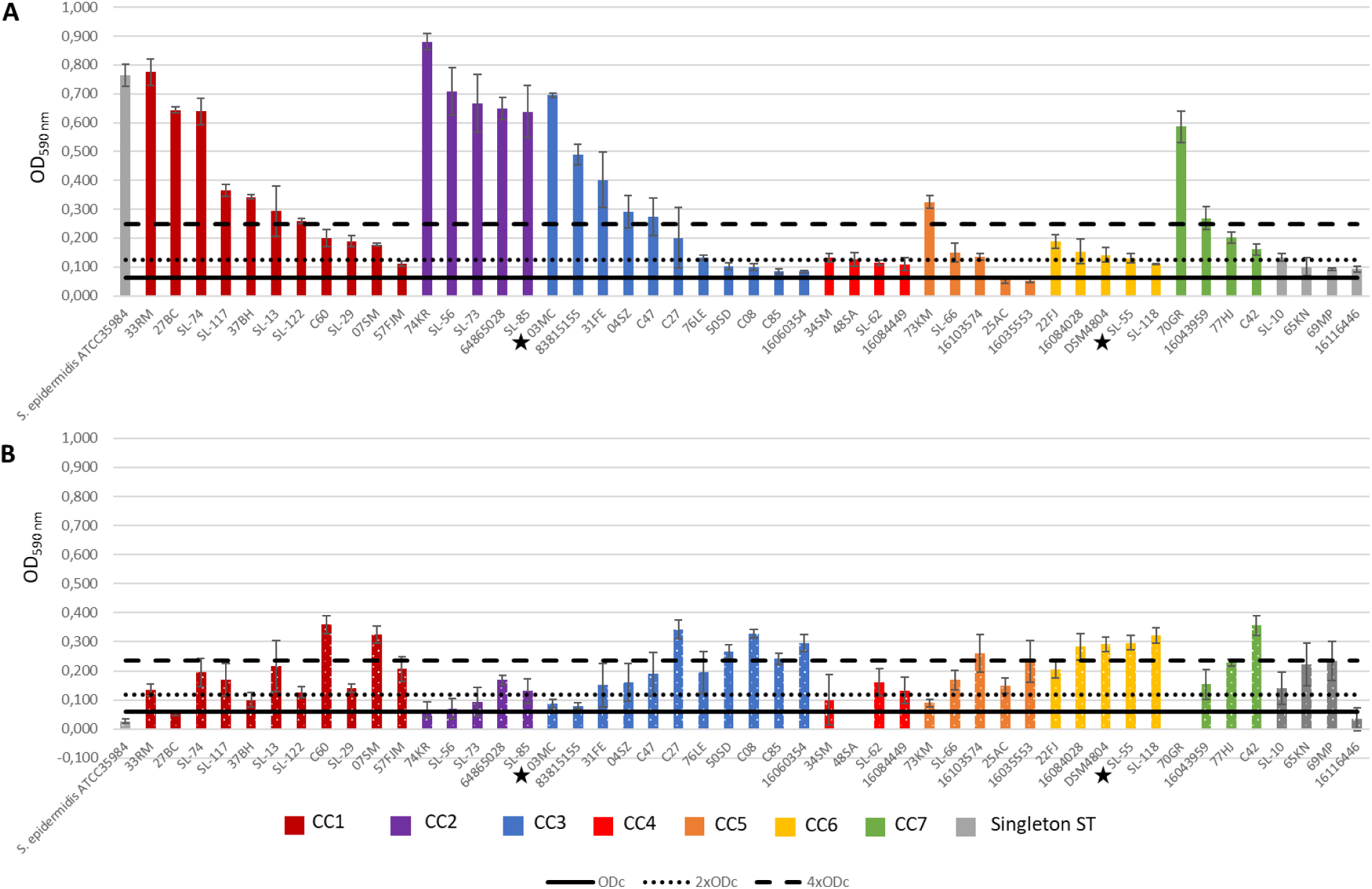
Biofilm formation of 49 *S. lugdunensis* strains in rich (TSBG) (A), or iron deficient medium (RPMI) (B). Biofilm formation was evaluated by crystal violet staining from four independent biological experiments. Strains were classified as strong-(4 x ODc < OD), moderate-(2 x ODc < OD ≤ 4 x ODc), weak-(ODc < OD ≤ 2 x ODc) and non-biofilm producers (OD ≤ ODc) according to the final OD value. ODc = Average OD_590_^blank^ + 3 ∗ Standard Deviation^blank^(25). Strains were classified by CC. The two strains marked with a star are those whose biofilm was further characterized.

**Figure 2.**
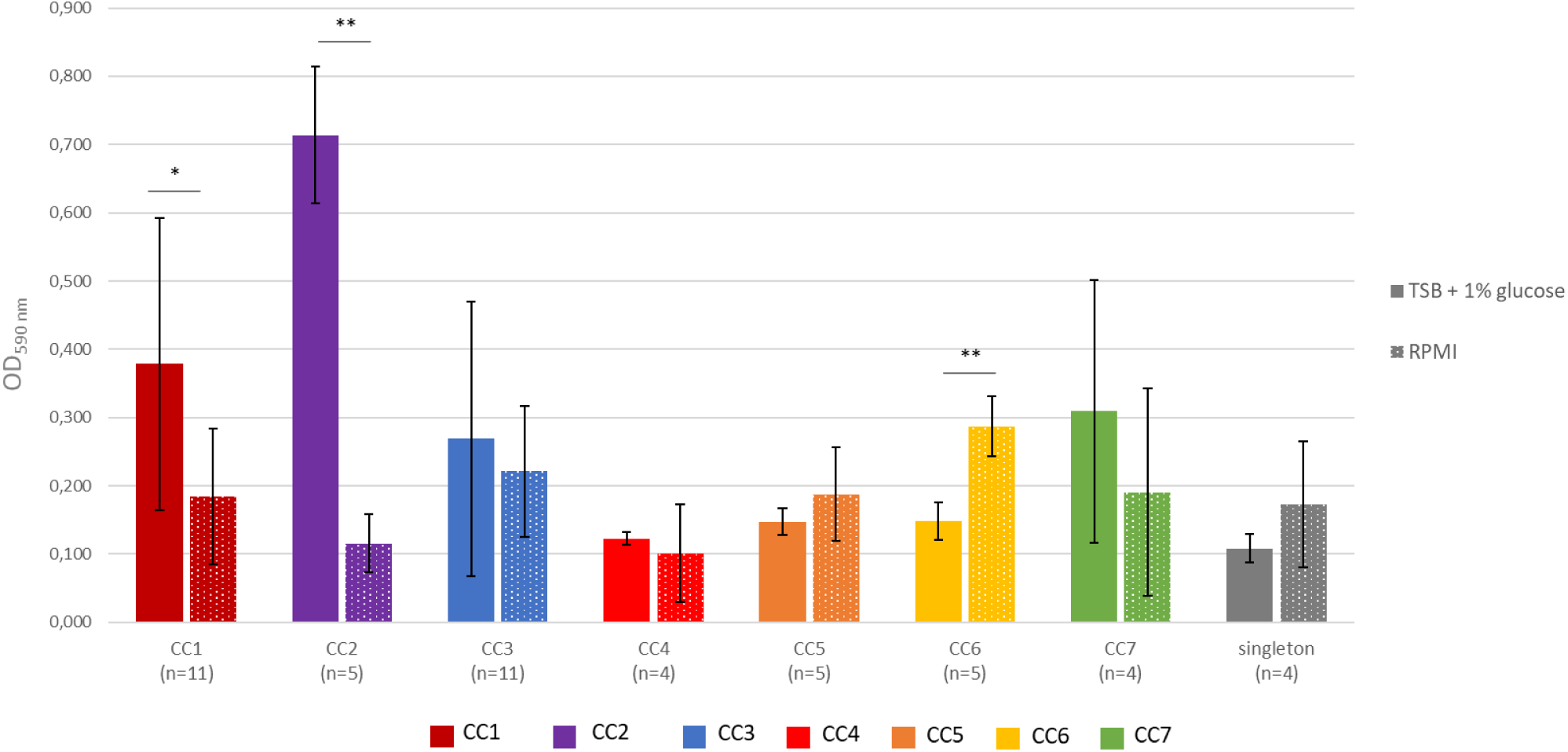
Biofilm formation comparison of 49 *S. lugdunensis* strains according to the CCs (OD_590_ means) in rich-(TSBG, full bars) and iron-restricted (RPMI, dotted bars) media. Error bars represent the standard deviation within each CC. Significance was identified as *p* < 0.05 (*) or *p* < 0.01 (**) by Kruskal–Wallis test and post hoc Dunn test. Data are the result of four independent biological experiments.

In iron-restricted conditions (RPMI), 45/49 strains (91.8%) produced biofilm. However, strains produced less biofilm than in TSBG (OD_590 max_= 0.359 ± 0.062 in RPMI *versus* 0.897 ± 0.072 in TSBG). 26.5% (*n* = 13) were strong-, 49% (*n* = 24) moderate- and 16.3% (*n* = 8) weak-biofilm producers (***Figure 1B***). Strains from CC6 formed significantly more biofilm than those from CC1 (*p* = 0.016), CC2 (*p* = 0.008), CC3 (*p* = 0.004), CC4 (*p* = 0.016) and CC5 (*p* = 0.032) (***Figure 2***).

Whatever the medium, the biofilm production was not correlated to the clinical context (infection *vs* carriage) or the presence of medical devices (***Supplementary Table 1***). However, when comparing biofilm formation in TSBG and RPMI, two CC-related behaviors were observed. On average, CC1 (*p* = 0.013) and CC2 (*p* = 0.0079) strains produced significantly more biofilm in rich medium than in iron-restricted conditions, while CC6 strains (*p* = 0.0079) were significantly more productive in RPMI (***Figure 2***). Of note, a heterogeneous behavior was observed among the CC3, with some strains being much higher producer (*e.g*. 03MC) in rich conditions (***Figure 1***).

### Biofilm composition and architecture differ between two *S. lugdunensis* strains of different CC

Two *S. lugdunensis* strains were selected for further biofilm composition analysis, due to their different behaviors: one CC3 strain (03MC, isolated from skin and soft tissue infection, strong-biofilm producer in rich medium and weak-producer in iron-restricted conditions) and one CC6 strain (SL-55, isolated from skin and soft tissue infection, moderate-producer in rich medium and strong-producer in iron-restricted conditions).

To investigate the role of eDNA, extracellular proteins, and polysaccharides in the first steps of biofilm formation, DNAse I, proteinase K and metaperiodate were used to alter each of these potential matrix constituents. Both strains showed a significantly (*p* < 0.05) reduced biofilm formation in presence of DNAse I and proteinase K when added at the initial point of inoculation in rich medium (TSBG) (***Figure 3***), the CC3 strain being more affected than the CC6 one. When using a pretreatment by metaperiodate, the CC3 strain biofilm formation was significantly reduced (*p* = 0.0049), which was not the case for the CC6 strain.

**Figure 3.**
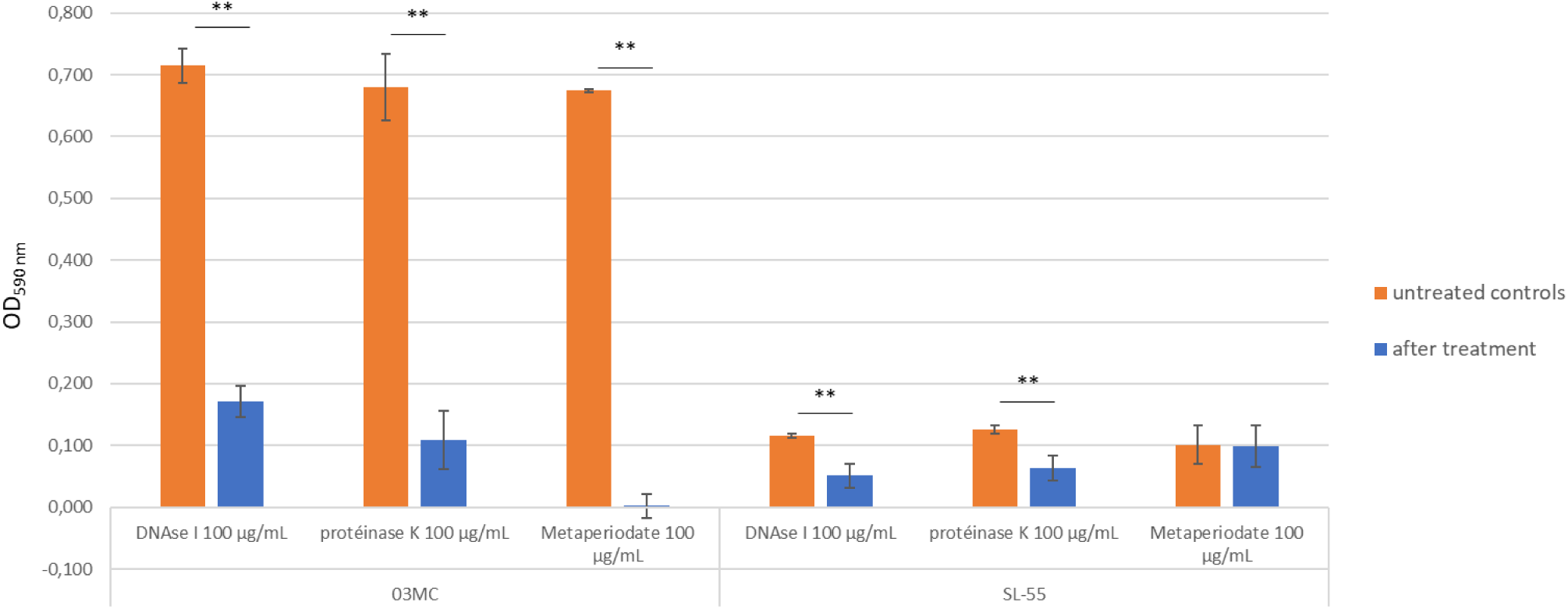
CC3 (03MC) and CC6 (SL-55) *S. lugdunensis* strains biofilm inhibition assays in presence of DNAse I (100µL/mL), proteinase K (100µL/mL) and metaperiodate (100µL/mL), from three independent biological assays. Error bars represent the standard deviation of stained biofilm. Significance was identified as *p* < 0.01 (**) by Wilcoxon Mann Whithney test.

The architecture, cell viability and ECM composition of a 48-h biofilm was further investigated using CLSM for both strains. In rich medium, CC3 and CC6 strains were able to form a flat and homogeneous biofilm, with a maximal thickness of 22,06 ± 0.63 µm and 26.85 ± 0.47, and an average thickness for live cells of 18.66 ± 0.41 µm and 15.06 ± 0.49 µm, respectively (***Supplementary figure 1***). Of note, CC3 strain biofilm was more compact than that of CC6 strain with roughness of 0.12 *versus* 0.32, respectively. Live/dead labeling showed that CC3 strain biofilm was mainly composed of viable cells, representing about 72.5% of the total cells, while the biofilm of CC6 strain was mainly composed of dead cells (61.6%) (***Figure 4***). Noticeably, in iron-restricted medium, CC6 strain displayed a biofilm with increased biovolumes, maximal and average thicknesses compared to CC3 strain. In this condition, the ratio of dead cells increased in CC3 strain biofilm, reaching 54.8% of the total population, while the rate of dead cells in CC6 strain biofilm (47%) decreased compared to TSBG (***Figure 4***).

**Figure 4.**
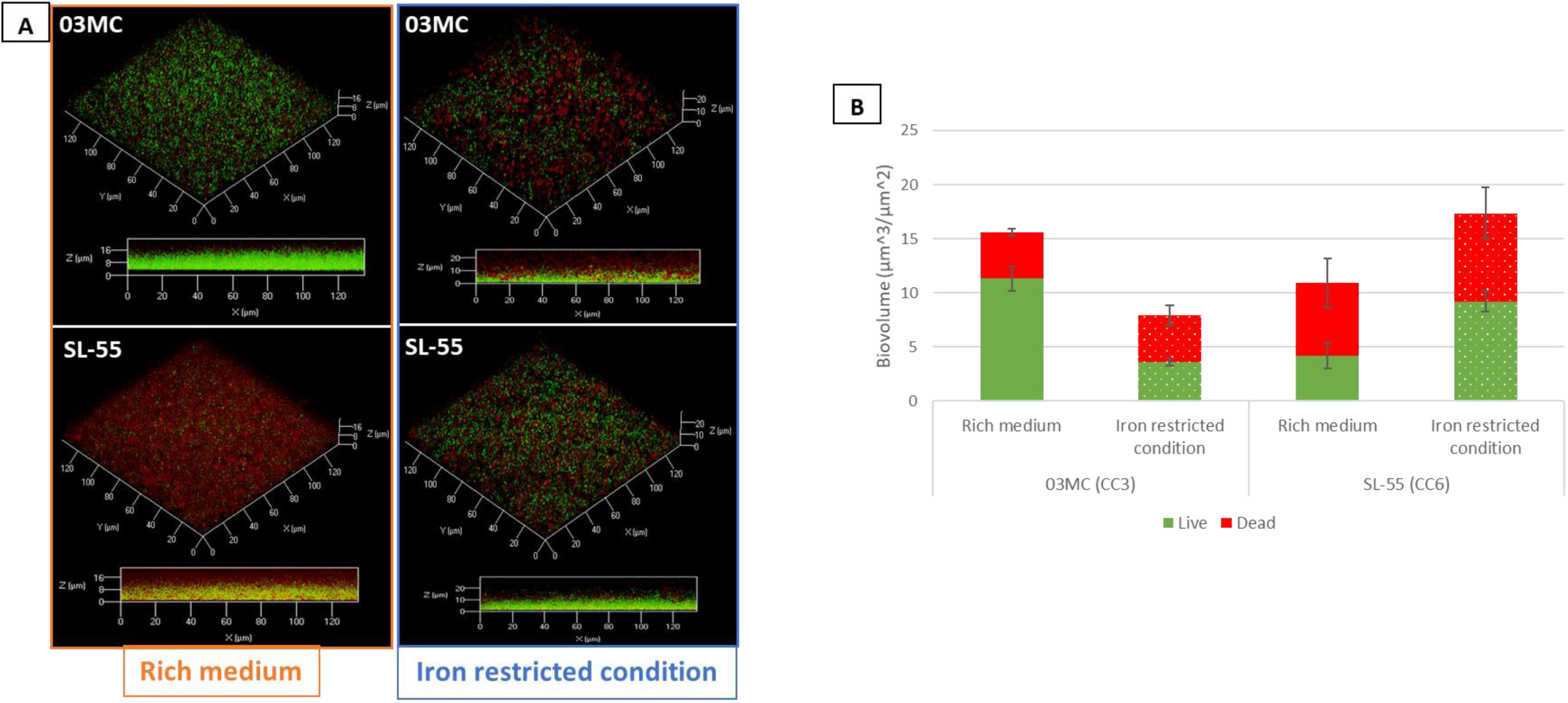
CC3 (03MC) and CC6 (SL-55) *S. lugdunensis* strains biofilm formation of in rich (TSBG) and iron restricted media (RPMI). 3D representations of biofilm assessed by cellular live/dead staining and observed by CLSM for CC3 (03MC) and CC6 (SL-55) strains in TSBG and RPMI (A). COMSTAT image analysis of biofilm composition of living (green) and dead (red) cells (B). Data are the results of the analysis of 10 views from three independent experiments.

Looking at the ECM of these biofilms, in rich medium, CC3 strain was composed of relatively similar amounts of proteins, eDNA and polysaccharides with average thickness of 2.01 ± 0.18 µm, 3.77 ± 0.56 µm, and 1.49 ± 0.39 µm respectively, while the ECM of the CC6 strain was predominantly proteinaceous (average thickness of 4.98 ± 0.30 µm) (***Figure 5***, ***Supplementary figure 1***). In iron-restricted, CC3 strain ECM was composed of proteins and polysaccharides in similar proportions (average thickness of 3.4 ± 0.2 µm and 2.27 ± 0.25 µm respectively) and eDNA in higher amounts (5.51 ± 0.4 µm), while CC6 strain contained a lower proportion of polysaccharides (1.01 ± 0.1 µm) than proteins (2.86 ± 0.3 µm) and eDNA (2.26 ± 0.13 µm). CC3 strain produced significantly larger amounts of ECM than in TSBG (*p*<0.05), with a lower proportion (not significant) of proteins than in TSBG. CC6 strain produced slightly lower amounts of ECM in RPMI, with a propensity for a lower proportion of proteins (***Figure 5***, ***Supplementary figure 1***).

**Figure 5.**
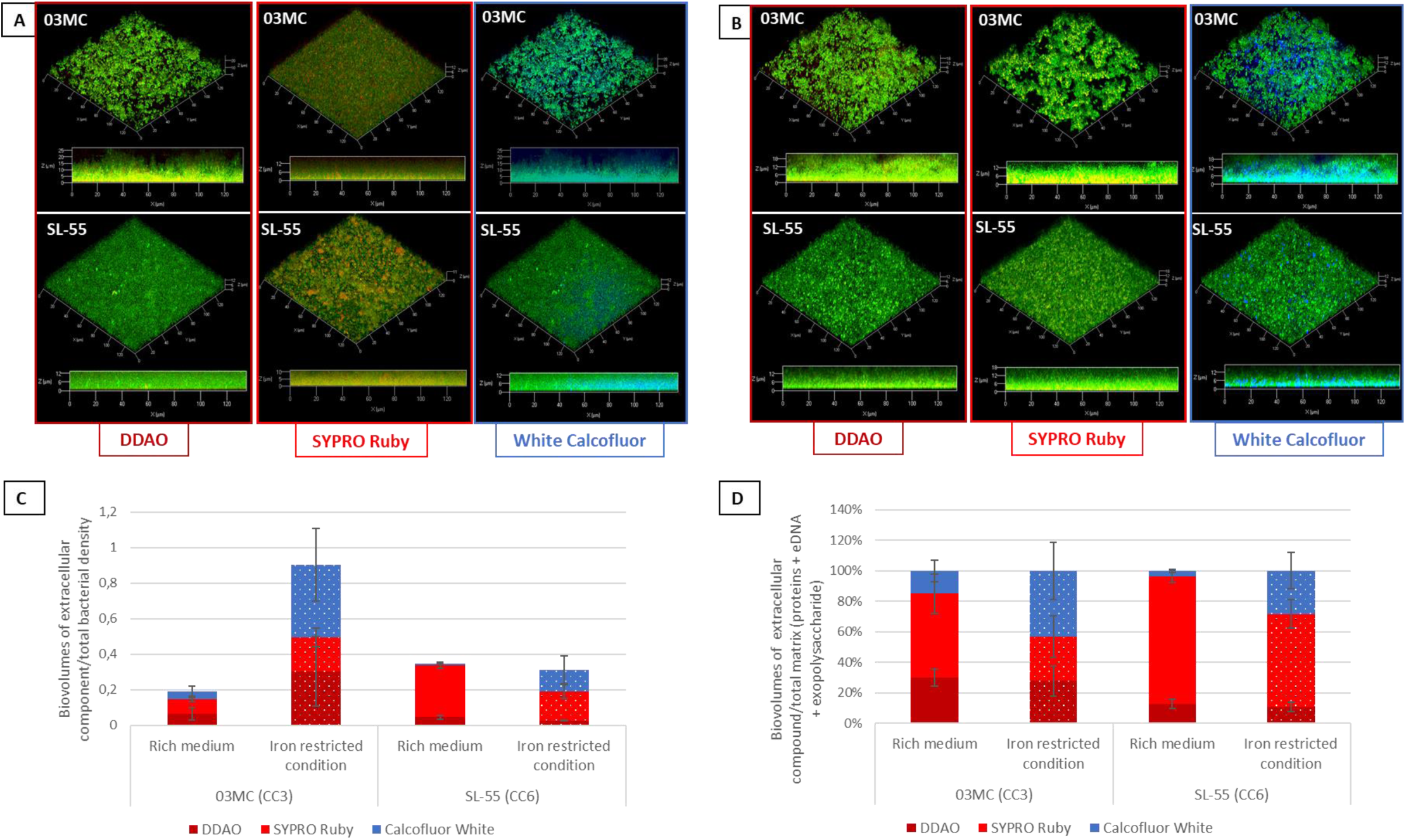
Extracellular matrix of *S. lugdunensis* biofilm. 3D representations of *S. lugdunensis* biofilm after labelling of live cells, proteins, eDNA, and polysaccharides by Syto9 (green), SYPRO ruby (red), DDAO (red) and Calcofluor White (blue) respectively in TSBG (A) and RPMI (B). COMSTAT image analysis of biofilm matrix composition (eDNA, proteins, and polysaccharides) relative to the total bacterial density (C), and relative to the total matrix (D). Data are the results of the analysis of 10 views from three independent experiments.

### In rich media, the matrix protein composition of the two strains differs in terms of abundance

ECM proteomes of the 24-h biofilm of CC3 and CC6 strains in TSBG were obtained after analysis of the matrix extracts by LC-MS/MS. A total of 321 proteins was identified (*i.e.* proteins identified from at least 2 peptides) (***Supplementary table 2***); no protein was specific to either strain. These proteins were mainly identified as intracellular (71.7%), membrane proteins (13.7%), and a few ones were extracellular (4%) (10.6% were of unknown location). Ribosomal proteins accounted for 12.1% (*n* = 39) of total proteins (***Table 1***). Among membrane proteins, eight were ABC transporters, six were two-component systems and four were LPXTG-containing surface proteins (SLGD_00094, IsdA; SLGD_0090, IsdB; IsdC; and SLGD_00478, a triacylglycerol lipase). In addition, 12 transcriptional regulators were identified, including two from the LytR family (SLGD_01820 and SLGD_00719). Of the 20 most abundant proteins, 11 were common to both strains (***Table 2***). These included a LTA synthase family protein (SLGD_02116, lipoteichoic acid synthase), metabolism proteins, a LysM peptidoglycan-binding domain-containing protein (SLGD_02450, N-acetylmuramoyl-L-alanine amidase) and an autolysin (SLGD_02076, N-acetylmuramoyl-L-alanine amidase). It is noteworthy that two proteins (SLGD_02212 and SLGD_00206) were highly abundant in both strains (abundance > 1,000,000). According to the NCBI database, SLGD_02212 corresponded to a SA0570 family protein, and SLGD_00206 to an immunodominant staphylococcal antigen IsaB family protein.

**Table 1.** Localization of proteins identified in the ECM matrix of *S. lugdunensis* CC3 and CC6 strains.

| | Total identified<br>proteins ($n = 321$) | Proteins significantly more<br>abundant in CC3 strain<br>($n = 148$) | Proteins significantly more<br>abundant in CC6 strain<br>($n = 54$) |
| --- | --- | --- | --- |
| Intracellular | 230 (71.7%) | 110 (74.3%) | 36 (66.7%) |
| Extracellular | 13 (4%) | 4 (2.7%) | 5 (9.3%) |
| Membrane | 44 (13.7%) | 25 (16.9%) | 3 (5.6%) |
| Unknown | 34 (10.6%) | 9 (6.1%) | 10 (18.5%) |

**Table 2.**
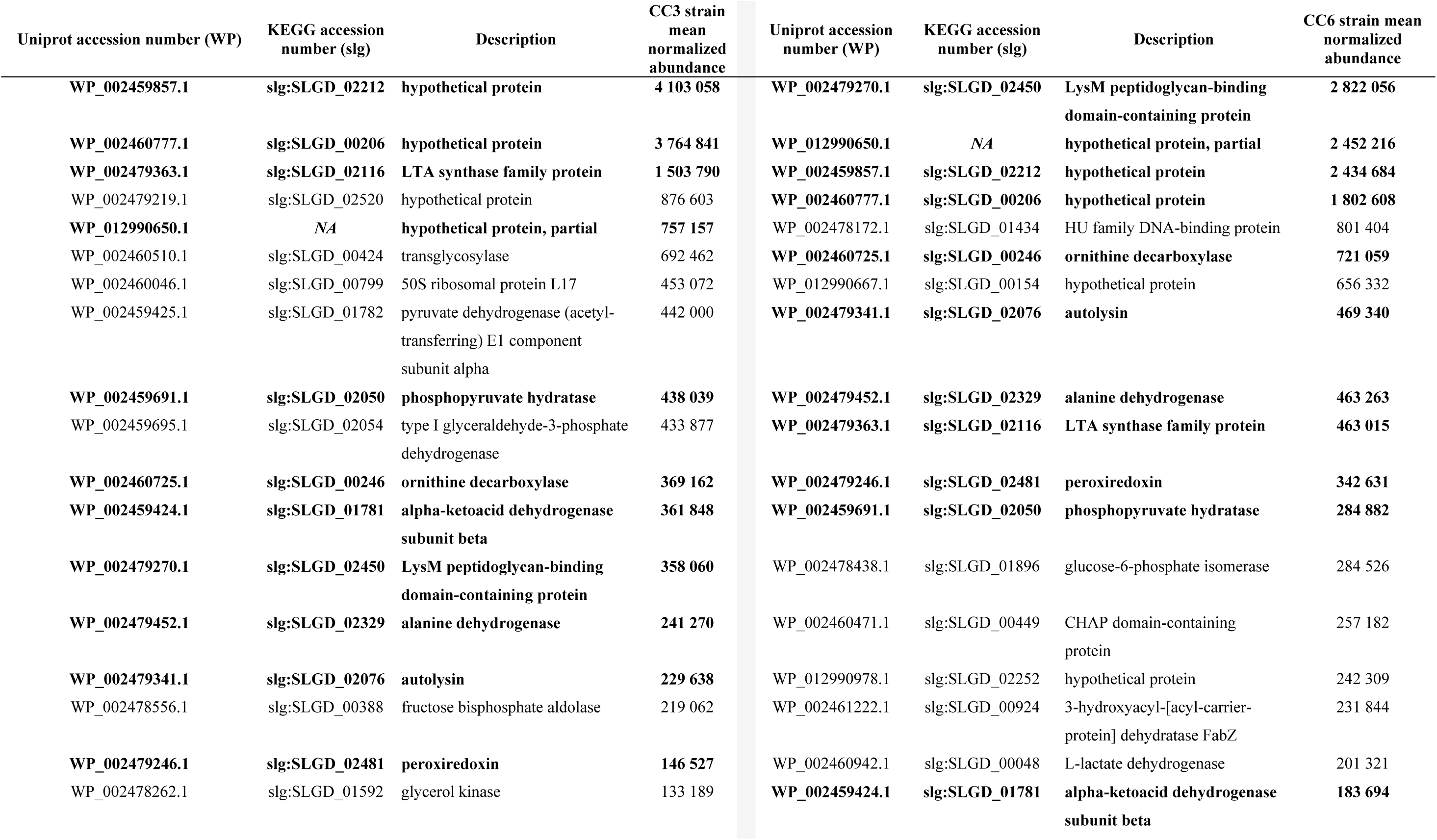
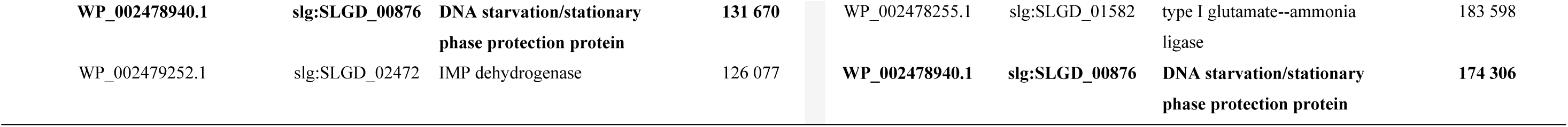
The 20 most abundant proteins among all identified proteins (*n* = 321) in the biofilm matrix of the CC3 and the CC6 strains in TSBG (list in order of abundance for each strain). Proteins common to both strains are in bold. NA: not applicable.

Functional pathways of the 321 proteins were obtained via KEGG (***Figure 6***). The main functional category was linked to metabolism (72% of proteins), followed by genetic information processing (22%) and environmental information processing (4%). The most represented pathway in the metabolism category was the carbohydrate metabolism (*n* = 98 proteins); in this pathway, proteins were strongly involved in glycolysis/gluconeogenesis (*n* = 22) and pyruvate metabolism (*n* = 14), but more rarely in citrate cycle and tricarboxylic acid (TCA) cycle (*n* = 9).

**Figure 6.**
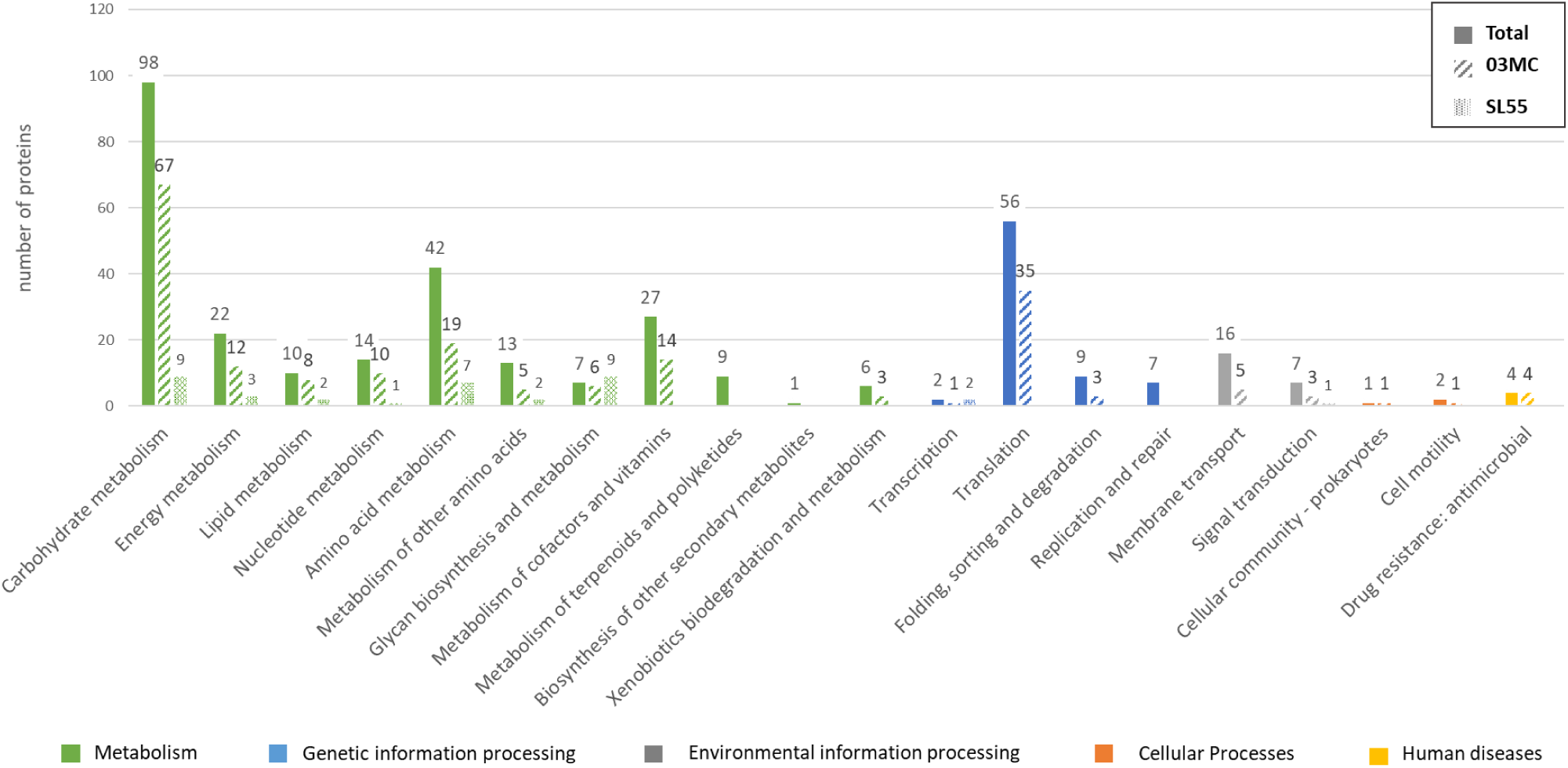
Functional classification according to KEGG pathways of the 321 proteins identified in the TSBG biofilm matrix of *S. lugdunensis* CC3 (03MC) and CC6 (SL-55) strains (full bars), and upregulated in CC3 strain (hatched bars) and CC6 strain (dotted bars). The number of proteins of each channel is indicated above each histogram bar.

Of the 321 proteins identified, 119 were considered to be non-varying between the two strains, as they had a *p*-value >0.05 and/or a power >0.8 and/or a fold change <1.8. Of the remaining proteins, 148 were significantly more abundant in the CC3 strain and 54 in the CC6 strain (***Supplementary table 2***). In both strains, the significantly more abundant proteins were mainly intracellular (74.3% and 66.7% for CC3 and CC6 strains respectively). We found a higher proportion of membrane proteins among the more abundant proteins in the CC3 strain (16.9%) than in the CC6 strain (5.6%) (***Table 1***); these included IsdB (SLGD_0090), three ABC transporters and two phosphotransferase systems (***Supplementary table 2***). Moreover, LytR (SLGD_01820) was present among the twenty most abundant overexpressed proteins found in CC3 strain (***Supplementary table 3***). The most represented pathway in both strains was metabolism (97.3% [144/148] and 61.1% [33/54] of proteins for CC3 and CC6 strains, respectively), and more specifically carbohydrate metabolism (***Figure 6***). A higher percentage of proteins involved in processing genetic information was observed for CC3 strain compared to CC6 strain (26.4% *vs* 3.7%, respectively), with most proteins (35/39) being involved in translation.

## DISCUSSION

Biofilm is strongly associated with persistence and difficult-to-treat infections (10, 30, 31). Concurrently, iron acquisition is critical for pathogenic bacteria to survive, colonize and invade host tissues. We therefore evaluated the impact of iron-restricted conditions (RPMI), compared to rich medium (TSBG), on bacterial growth and biofilm-forming capacity of a large selection of *S. lugdunensis* clinical strains belonging to the seven CCs defined by MLST to date.

This study reveals that *S. lugdunensis* is a biofilm-producing species in both iron-rich and iron-poor environments, with production levels varying among strains. In TSBG, our results are consistent with those of previous studies using a microtiter plate assay, such as Frank and Patel (13), Qian *et al*. (15), Missineo *et al.*(21), and Hagstrand Aldman *et al.* (10) studies in which all strains were biofilm producers, but greatly higher than the findings of Pereira *et al.* (60.9% producers) (17). In iron-restricted medium, our strains formed on average significantly less biofilm than in rich medium, probably in connection with a reduced growth rate in RPMI. However, some strains, particularly CC6 strains, formed significantly more biofilm in this *in vivo* like condition. This result is in link with the findings of Missineo *et al.* where their nine studied strains (unknown CC) were biofilm producers (100%) in similar conditions (RPMI with 0.3% glucose and 2 mM glutamine) (21). In the same way, Aubourg *et al.* reported that iron limitation promoted biofilm formation of the strain *S. lugdunensis* N920143, but this was obtained in different conditions (brain-heart infusion broth with 2,2’-dipyridyl as iron chelator) and on a single strain (20). Moreover, this study shows for the first time that the biofilm formation of *S. lugdunensis* is associated with the phylogenetic lineage, with two CC-dependent behaviors (CC1 and CC2 strains producing significantly more biofilm in rich medium than in iron-restricted medium, while the opposite was observed for CC6 strains). It is noteworthy that the high-prevalent clones CC1 and CC3 were, on average, strong-biofilm producers, but exhibited strain-to-strain heterogeneity. Similarly, for *S. aureus*, a correlation between biofilm phenotype and clonal lineages defined by MLST and/or *spa* typing has been described (32–34).

We further investigated the composition and the structure of *S. lugdunensis* biofilm by selecting two strains that best represented these types of behaviors: one strain belonging to the high-prevalent clone CC3 (03MC, strong-biofilm producer in rich medium) and one CC6 strain (SL-55, strong biofilm producer in RPMI). The biofilm formation and composition varied according to the strain and culture medium, as previously pointed out (15, 16). Biofilm inhibition assays results indicated that in rich media, both proteins and eDNA play key role in the development and the structural integrity of *S. lugdunensis* biofilms, while the role of polysaccharides is strain-dependent. Indeed, the matrix of the CC6 strain was essentially proteinaceous without exopolysaccharides, which could be linked to a frameshift of *icaA* gene resulting in an early stop in IcaA (involved in synthesis of N-acetylglucosamine oligomers, 114 amino acids instead of 407) (https://www.ncbi.nlm.nih.gov/datasets/genome/GCF_008728735.1/). Data obtained by CLSM showed that the two strains of *S. lugdunensis* of different clonal lineage presented different biofilms in terms of architecture and cell viability, depending on their growth medium. Notably, in iron-restricted conditions, the biofilm of the CC3 strain contained more dead cells, which could be related to higher eDNA level, while that of the CC6 strain contained more living cells (35, 36). Second, the ECM of both strains contained proportionally less proteins but more polysaccharides than in rich medium, suggesting that polysaccharides would play a more relevant role in the structural integrity of the biofilm in RPMI, contrary to what was shown by Missineo *et al*. (21). However, it should be noted that in our study exopolysaccharides were stained using Calcofluor White (as described by Grecka *et al*. [(37)]), which marks β-1,3 and β-1,4 polysaccharide bonds, whereas staphylococcal ECM has been described to date as mainly composed of PNAG which contains β-1-6 linkages (38). Thus, this suggests that polysaccharides other than PNAG could compose the biofilm matrix of *S. lugdunensis* strains (39). Further studies should consider completing these CLSM analyses by using a marker targeting the beta 1-6 bonds of PNAG, such as wheat germ agglutinin (13).

Studying the protein composition of the matrix is of importance for understanding bacterial lifestyle and developing new therapeutic options against biofilm-associated infections. Only one study recently investigated the protein composition of the biofilm matrix of one *S. lugdunensis* strain (not phylogenetically characterized) (22). In the present study, biofilm formation screening of phylogenetically characterized strains enabled us to select two strains based on their CC (CC3 and CC6) as well as their biofilm formation ability. Analysis of the ECM proteome of these two representative strains has identified a total of 321 proteins, all common to both strains, which is low compared to the 1,125 identified by Cho *et al*. (22). This result may be related to differences in biofilm growth conditions (24-h biofilm in TSBG in our study versus 72-h biofilm in TSB without glucose in Cho *et al*’s study) as well as the extraction method used (chemical in our study and mechanical in Cho *et al*.’s study) (22). The vast majority of the proteins identified in our study were cytoplasmic (71%), similar to studies that characterized biofilm ECM of *S. aureus* and *S. epidermidis* strains (60% to 86.2% of intracellular proteins) (40–42). These intracellular proteins may originate (i) from cell lysis within the biofilm, which may be autolysis-dependant (in line with the identification with one extracellular autolysin in the ECM of our strains) (43), (ii) from non-classical protein export by an yet unknown pathway or (iii) from mechanical cell lysis during ECM extraction. Of the intracellular proteins identified here, 12.4% were ribosomal proteins, whereas this rate reached 27% in the matrix of *S. epidermidis* in Martinez *et al*. study (41). The accumulation of cytoplasmic proteins could correspond to a “moonlighting” strategy of bacteria to stabilize the biofilm structure (40, 44). Moreover, in our study membrane proteins accounted for 13% of the proteins identified in the matrix biofilm of *S. lugdunensis*. Among these proteins, eight were ABC transporters, and three were LPXTG-containing surface proteins, including three Isd proteins (IsdA, IsdB and IsdC). This result is in agreement with Cho *et al*.’s findings, who identified ABC transporters, one LPXTG-containing surface protein and the probable heme-iron transport system permease protein IsdF among the most abundant biofilm proteins, suggesting their role in the *S. lugdunensis* ECM composition (22). Identification of Isd proteins under rich conditions suggests that their expression is not limited to iron-limiting medium, unlike what was initially shown for IsdC (20, 21). Of note, among the 20 most abundant matrix proteins identified in the biofilm of both strains, was a LysM peptidoglycan-binding domain-containing protein which, in *S. aureus*, mediates adherence to the extracellular matrix (45). Interestingly, an LTA synthase was also found, essential for the synthesis of teichoic acids, which is important for *S. aureus* or *S. epidermidis* initial adhesion to biotic surfaces (12). Two proteins were identified as highly abundant in the matrix of both strains, but were classified as hypothetical proteins. These proteins have been described in *S. aureus*: one (SLGD_02212) is a protease (Clp) (46), this type of enzyme being involved in biofilm dispersal phase (12), and the other (SLGD_00206) is an IsaB analog protein, an extracellular nucleic acid binding protein whose role in virulence is not established in *S. aureus* (46). KEGG analyzes revealed that the biofilm matrix of the two *S. lugdunensis* strains mostly contained proteins involved in metabolic pathways, particularly in carbohydrate metabolism, as previously described in *S. lugdunensis* and other staphylococci (22, 41, 47, 48). In particular, we have identified a large number of proteins involved in glycolysis and gluconeogenesis for the formation of pyruvate, in line with the results obtained for *S. epidermidis* (41) or *S. aureus* biofilms (48). Similarly to *S. epidermidis* (41), few TCA cycle proteins were identified in our study in the ECM of *S. lugdunensis* biofilm, unlike *S. aureus* whose biofilm formation was initiated by metabolites produced by the TCA cycle (48). It would be interesting to compare proteomes of planktonic *versus* biofilm cultures of various strains of *S. lugdunensis* and to carry out metabolomic studies using metabolic techniques such as nuclear magnetic resonance (48) to confirm the involvement of these metabolic pathways biofilm formation in this species.

Interestingly, ECM of the two *S. lugdunensis* strains differed in terms of proteins abundance. The higher number of proteins more abundant in CC3 strain compared to CC6 strain (148 *vs*. 54), including a higher number of membrane proteins, may be related to its strong ability to produce biofilm under rich conditions, as membrane proteins are involved in the early stages of biofilm formation (49, 50). Genetic information processing proteins were more abundant in CC3 strain than in CC6 strain and included 22 ribosomal proteins, supporting the role of these proteins in biofilm formation. Furthermore, LytR, which was more abundant in CC3 strain biofilm, is a transcriptional regulator belonging to the two-component LytSR regulatory system, previously shown to be involved in biofilm formation in *S. lugdunensis* (51, 52).

In conclusion, our study is the first to investigate the biofilm-forming capacity of such a large collection of *S. lugdunensis* strains, representative of all the CCs, in rich- and iron-restricted media. We demonstrated for the first time that biofilm formation was strongly associated with CC affiliation and iron availability. Furthermore, the in-depth biofilm characterization of two strains belonging to two CCs using complementary approaches, including proteomic characterization of the ECM, confirmed notable differences in biofilm composition.

## ACKNOWLEDGMENTS

The authors would like to thank the PISSARO Platform HERACLES US 51 UAR 2026 of Rouen Normandy University for LC-MS/MS analyses

## CONFLICTS OF INTEREST

The authors declare that the research was conducted in the absence of any commercial or financial relationships that could be construed as a potential conflict of interest.

